# Quantitative mapping of methionine sensitivity to oxidation in the copper-bound PcuC chaperone

**DOI:** 10.1101/2025.09.11.675508

**Authors:** Lionel Tarrago, Lise Molinelli, Maya Belghazi, Mathilde Tribout, David Lemaire, Pierre Legrand, Sandrine Grosse, David Pignol, Monique Sabaty, Thierry Tron, Pascal Arnoux

## Abstract

Copper is typically coordinated by histidine, cysteine, or methionine in proteins, and these residues are particularly sensitive to oxidation. However, it remains unclear whether copper-coordinating residues are more prone to oxidation than non-coordinating ones, and how their susceptibility changes between the apo and copper-bound states. The copper chaperone PcuC, important for cytochrome c oxidase assembly in bacteria, contains a canonical binding site composed of two histidines and two methionines (H51x*_n_*M63x*_22_*H86xM88), as well as a disordered C-terminal extension enriched in methionine and histidine. To quantify methionine oxidation sensitivity in both apo- and Cu-bound PcuC, we used a methionine-specific oxaziridine probe combined with mass spectrometry and compared labeling patterns to those generated by ^18^O-labeled hydrogen peroxide. We show that methionine residues display distinct oxidation sensitivities in the apoprotein, and that the oxaziridine reacts similarly to H_2_^18^O_2_. Importantly, this probe enables quantification of methionine oxidation independently of hydroxyl radicals generated by copper-driven Fenton chemistry, which lacks residue specificity. In the copper-bound form, Cu binding strongly alters methionine reactivity, with a marked increase in oxidation of the coordinating Met63 and Met88. Structural analysis revealed that two copper ions occupy the canonical site, while the C-terminal extension does not contribute to coordination. Comparison of structural features and oxidation values showed that methionine sensitivity correlates with solvent exposure in the folded domain, but with local positive charge in the disordered region. These findings demonstrate that copper coordination modulates methionine oxidation, and that oxaziridine-based probes provide powerful tools for mapping oxidation sensitivity in (metallo)proteins.

## 1. Introduction

Metals are essential in life, especially within proteins where they serve as critical cofactors for various biochemical reactions. Copper plays essential roles in biology as redox center in proteins involved in electron transfer, redox catalysis, and metal homeostasis [1]. Copper is commonly coordinated by histidine (His), cysteine (Cys), and methionine (Met) residues, with His and Cys predominantly found in redox-active sites while Met is mostly found in copper transporters and chaperone [2]. Copper can also bind proteins implicated in neurodegenerative diseases, such as α-synuclein and the prion protein [3,4]. The diversity of copper sites and ligands reflects the metal’s versatility, but also the need for precise control to ensure correct metalation and prevent oxidative damage. Indeed, the copper-coordinating residues Cys, His, and Met are particularly sensitive to oxidation, leading to radical or oxygenated forms. Oxidants like hydrogen peroxide (H_2_O_2_) or hypochlorite have distinct reactivity towards amino acids, and the reaction of H_2_O_2_ with iron or copper in Fenton(-like) reaction generates strongly oxidant hydroxyl radicals [5,6]. While hypochlorite and hydroxyl radicals have broad reactivity and affect most amino acids and the protein backbone, H_2_O_2_ reacts specifically with Cys and Met [7]. In particular, Met can be oxidized into sulfoxide (MetO) and sulfone (MetO_2_) [7,8]. Protein oxidation may have diverse consequences, from damage to regulated changes in function or activity. However, little is known about the sensitivity to oxidation of Cys, His, or Met within copper-binding proteins. It is not known whether these residues that are present in copper coordination sites, either in the apo- or copper-bound state, present a susceptibility to oxidation different from non-coordinating counterparts. Understanding the sensitivity of copper proteins to oxidation is crucial, as it can lead to their inactivation, as observed for superoxide dismutase and lytic polysaccharide monooxygenases [9–12]. It can also affect the structural properties of α-synuclein and the prion protein, with potential pathological outcomes [13,14]. In peptides mimicking copper-binding sites of the eukaryotic Ctr1 copper transporter, few evidence indicates that Cys is highly sensitive to oxidation contrary to Met or His [15,16]. Oxidation sensitivity of copper-binding Met in α-synuclein has also been observed, though quantitative data are lacking. In bacteria, such mechanisms remain poorly understood, despite their particular relevance for periplasmic proteins, which are exposed to copper-induced hydroxyl radical production and rely on detoxification and export systems [5,17,18]. Consistently, it has been shown that the expression of the methionine sulfoxide reductase (Msr) P/Q system, that repairs periplasmic protein by reducing oxidized Met [19,20], is induced by the presence of an excess of copper in the periplasm through the activation of the CusSR two-component system [21]. Bacterial membranes and mitochondria harbor the respiratory chain, where cytochrome *c* oxidase (Cox) contains two copper atoms forming the CuA center within its periplasmic catalytic subunit CoxB, which is essential for oxygen reduction [22]. Bacterial CoxB requires two chaperones for its assembly: PrrC, homologous to eukaryotic Sco proteins, and PcuC, a cupredoxin-like protein. This protein is often referred to as PCu_A_C but we use PcuC in this article to follow uniform nomenclature of bacterial proteins. The sequence of events allowing Cu insertion in CoxB has been evaluated in several bacterial species [23–26]. In the model proposed for *Cereibacter sphaeroides*, PcuC acquires copper from the periplasm and transfers it to PrrC, which then assembles the Cu_A_ center with one Cu(I) and one Cu(II) [26]. Cu ions are sequentially transferred from one coordinating site to another [27,28]. Particularly, CoxB coordinates the two Cu with two Cys, two His, one Met and one Glu in a H*x_34_*C*x*E*x*C*x_3_*H*x_2_*M motif [24,29]. PrrC uses two Cys and one His in the C*x*_3_C*x*_n_H motif to coordinate a Cu(II) ion [24]. It has been shown that copper-coordinating Cys of both CoxB and PrrC can form disulfide bonds that need to be reduced by the dedicated thiol-reductase TplA before copper insertion [30]. PcuC is widely conserved among bacteria and most homologs possess a Cu(I)-binding motif composed of two Met and two His (Hx*_n_*Mx*_22_*HxM) and have an additional C-terminal extension rich in Met and His, proposed to bind a second copper ion [23,24,31]. Other types lack this extension or bind copper in a His-brace [24,32]. The *C. sphaeroides* PcuC is a representative model for studying copper’s role in protein oxidation.

In this work, we evaluated the sensitivity of PcuC’s Met to oxidation, either in the apo or in the copper-loaded protein by using two molecules reacting with Met: i) ^18^O-labeled hydrogen peroxide, and ii) an oxaziridine probe (**Fig. 1A**). We quantified the level of oxidation of each Met by trypsin proteolysis and liquid chromatography coupled with tandem mass spectrometry (LC-MS/MS). H_2_O_2_ is a natural oxidant potentially encountered by bacteria. In the absence of Cys, as it is the case for PcuC, it allows the specific oxidation of Met [7,20]. The use of ^18^O-labeling avoid the quantification of artifactual Met oxidation potentially arising in proteomic analysis [33–35]. However, H_2_O_2_ can react with the metal in the copper-loaded protein, generating highly reactive hydroxyl radicals that may oxidize peptide bonds and residues other than Met. To specifically quantify methionine sensitivity to oxidation in the copper-loaded protein independently of Fenton reaction, we used an oxaziridine probe designed for selective reactivity with Met [36,37]. This probe operates via nitrogen atom transfer, where the electrophilic nitrogen of the strained N–O bond reacts with the thioether sulfur, forming a stable sulfimide adduct (**Fig. 1B**). We first assessed Met sensitivity in the apo form using H_2_^18^O_2_ and then with the oxaziridine probe, allowing comparison of both approaches. Then, we applied the probe to the copper-loaded protein to evaluate the impact of Cu coordination on Met oxidation. Finally, we determined the three-dimensional structures of both apo and copper-bound PcuC and integrated these structural insights with oxidation data to identify determinants of Met susceptibility to oxidation.

**Fig. 1.**
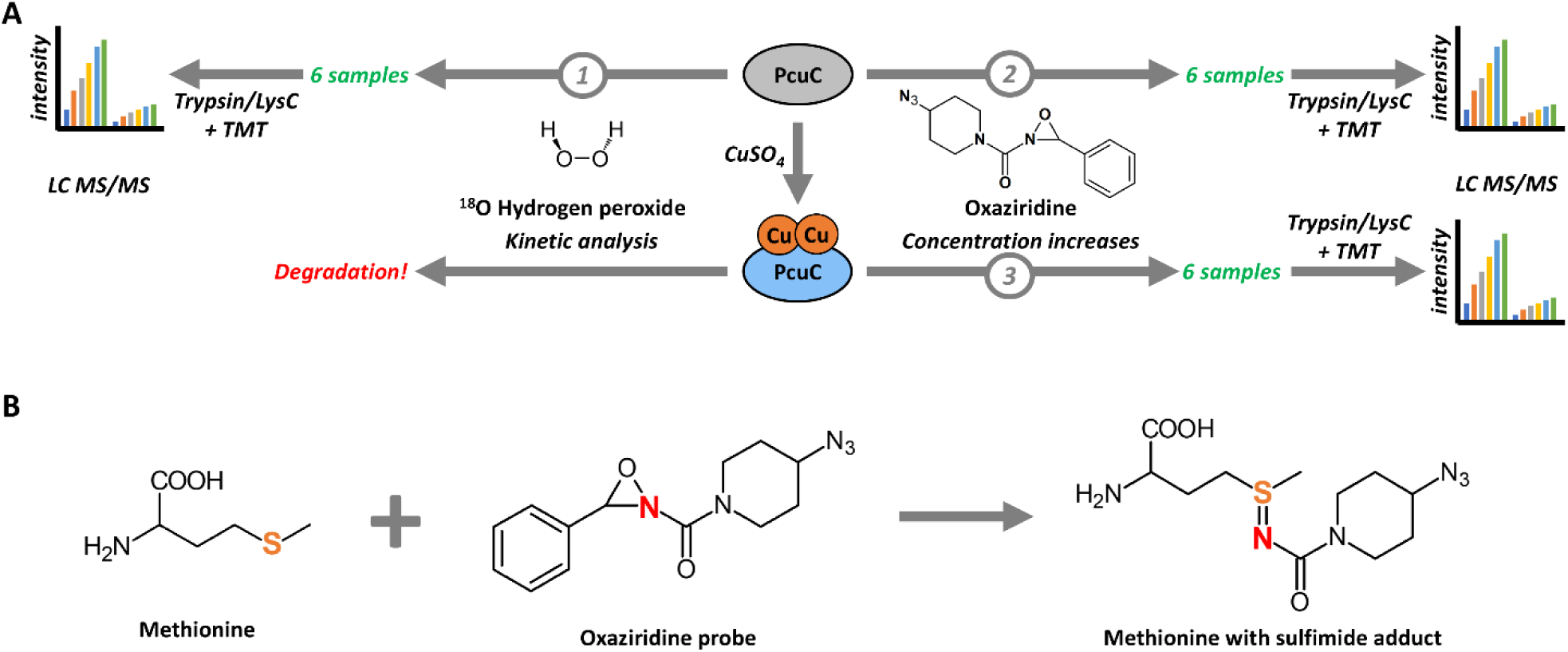
Utilization of an oxaziridine probe to quantify Met sensitivity to oxidation in PcuC and comparison with H_2_^18^O_2_. **(A)** Experimental design. Apo-PcuC was oxidized with H_2_^18^O_2_ in kinetic experiment (*1*) and apo-PcuC (*2*) or copper-bound PcuC (*3*) was incubated with increasing concentrations of oxaziridine probe. For these three analyses, six samples, prepared in triplicates, were digested with Trypsin/Lys C mix and analyzed simultaneously by LC-MS/MS using tandem mass tags (TMT) labeling. Analysis of Cu-loaded PcuC with H_2_^18^O_2_ was not performed because of protein degradation (*see* **Fig. S4B**). **(B)** The oxaziridine probe ((4-azidopiperidin-1-yl)(3-phenyl-1,2-oxaziridin-2-yl)methanone) oxidizes the sulfur atom in Met, resulting in a nitrogen transfer reaction. The sulfur of Met is covalently linked to the nitrogen atom of the oxaziridine probe via a sulfimide bond.

## 2. Results and Discussion

### 2.1. Methionine residues of the PcuC apoprotein have various sensitivity to oxidation

The copper chaperone PcuC contains eight Met residues: one isolated in the first half of the primary sequence (Met25), two pairs clustered together (Met61 and Met63, and Met88 and Met90), and three near the C-terminus (Met128, Met132, and Met135) (**Fig. 2A**). Using the purified recombinant apo-PcuC we first confirmed that it contains oxidizable Met residues and thus is relevant to quantify susceptibility to Met oxidation. It has up to seven MetO formed simultaneously by oxidation with H_2_O_2_ and most of which are reducible by the methionine sulfoxide reductase MsrP (**Supplemental Results 2.1, Fig. S1**). To quantify the sensitivity of the Met residues in the apo-PcuC individually, we incubated the protein with 1,000 molar equivalents of H_2_^18^O_2_ for 2 to 60 minutes and oxidized residue were quantified (**Fig. 2B, S2B**). An SDS-PAGE analysis of the samples revealed a delay in protein migration, as observed for non-isotopic H_2_O_2_, confirming that oxidation occurred (**Fig. S2A**). The identification and quantification of ^18^O-Met revealed that the most abundant form was ^18^O-sulfoxide, with only traces of ^18^O-containing sulfone detected (**Fig. S2B**). The search for oxidized form of His (oxoHis) revealed only extremely low levels, confirming that Met residues were the target of oxidation by H_2_^18^O_2_ (**Fig. S2C**). The analysis of the kinetics of oxidation revealed striking differences in the reactivity of the Met residues (**Fig. 2B, E**). Indeed, Met25 remained reduced throughout the treatment while Met135 was rapidly and extensively oxidized, up to ∼93% after 60 minutes. Met63, which is part of the copper-coordinating site, was the second most oxidized residue, reaching ∼45% by the end of the kinetics. The other Met residues showed intermediate levels of oxidation, all remaining below 50%. Interestingly, the oxidation kinetics of Met132 displayed a unique pattern compared to the other reactive Met residues. While for the other residues oxidation level quickly reached a plateau or increased linearly (Met61, Met128), Met132 oxidation level remained low at early timepoints but eventually increased to ∼42%, like the level observed for Met63 (**Fig. 2B**). This behavior suggests that Met132 may be a secondary target for oxidation, becoming oxidized only after the more sensitive residues, Met63, Met90, and Met135. Given the proximity of Met132 and Met135 within the disordered C-terminal region, preferential oxidation of the highly reactive Met135 may transiently limit oxidation of Met132. Based on final oxidation levels, Met reactivity in apo-PcuC ranked from least to most oxidized: Met25, Met128, Met88, Met61, Met90, Met132, Met63, Met135.

**Fig. 2.**
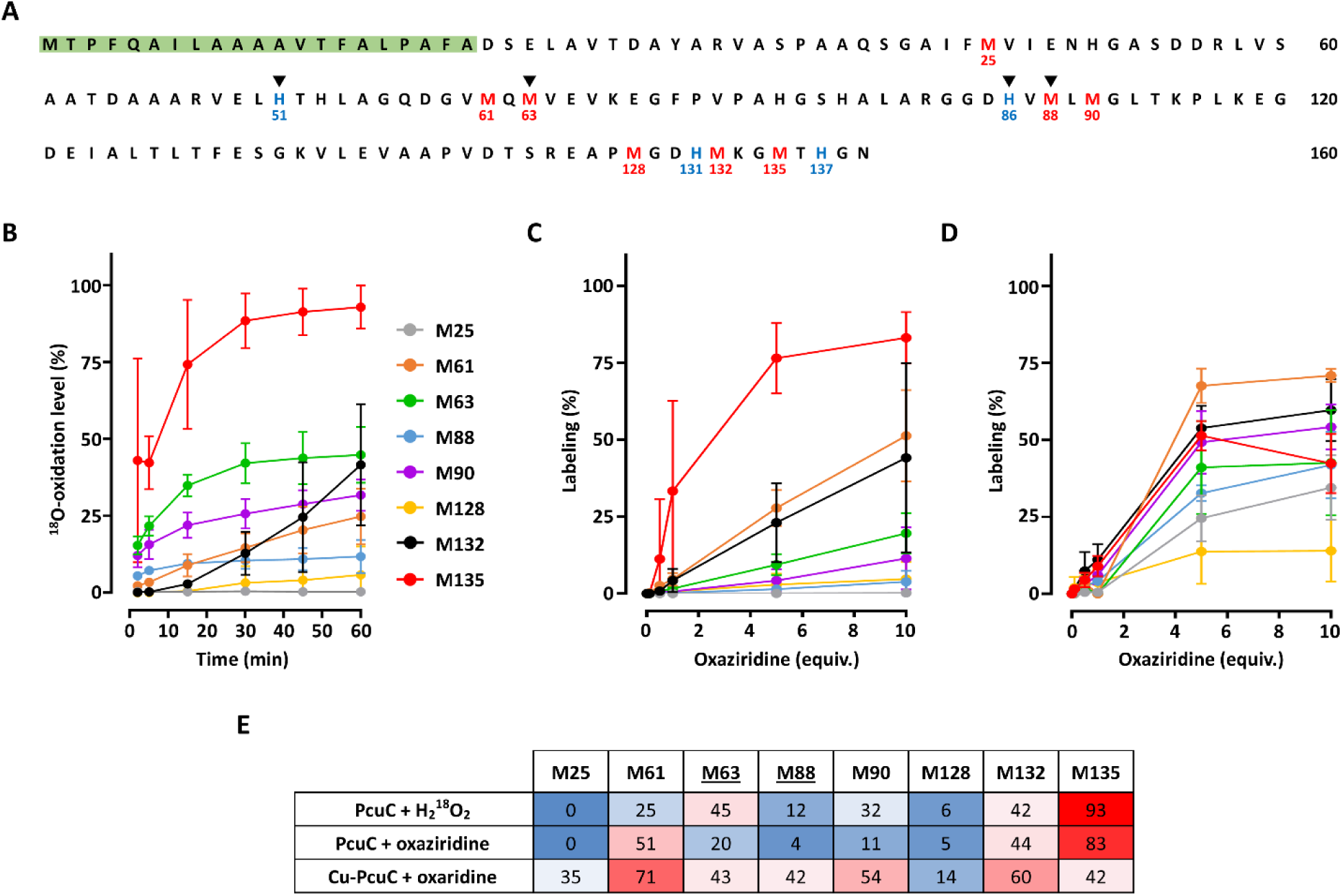
Quantification of Met sensitivity to oxidation in apo- and Cu-loaded PcuC using H_2_ ^18^O_2_ and an oxaziridine probe. **(A)** Amino acid sequence of PcuC in single letter. The eight Met residues are shown in *red* with numbering below, amino acids on *green* background correspond to the periplasmic peptide signal and *black* triangles indicate Cu-coordinating residues. Met are numbered considering Asp as first residue after removal of the peptide signal. **(B)** Time-resolved percentage of oxidation of the eight Met residues of the apo-PcuC after incubation with H_2_^18^O_2_. Apo-PcuC was incubated with 1,000 molar equivalents (equiv.) of H_2_^18^O_2_ for 2, 5, 15, 30, 45 and 60 min and the reaction stopped with catalase. Percentages were calculated by summing the relative intensities of all Met forms, grouping ¹□O-containing species as oxidized and others as non-oxidized. **(C)** Percentage of labeling of the apo-PcuC after incubation with the oxaziridine probe. Apo-PcuC was incubated with 0, 0.1, 0.5, 1, 5 or 10 equiv. of the oxaziridine probe and the reaction stopped by desalting. **(D)** Percentage of labeling of the Cu-loaded PcuC after incubation with the oxaziridine probe. Cu-loaded PcuC was incubated with the oxaziridine probe similarly to the apoprotein. Note that for Met135, the lower mean value at 10 versus 5 equivalents falls within the experimental standard deviation. **(E)** Heatmap showing the final percentage of oxidation or labeling of the eight Met residues of PcuC, using a *blue*-to-*red* color scale, with *blue* and *red* representing the lowest and highest values of modification, respectively. Underlined Met63 and Met88 belong to the Cu-binding site.

### 2.2. The oxaziridine probe reactivity toward Met is similar to that of hydrogen peroxide

With the aim of quantifying Met sensitivity in the copper-loaded PcuC, we used an oxaziridine probe [36,37]. First, we validated its ability to bind the apo-PcuC and its potential usability to quantify the sensitivity of PcuC’s Met residues to oxidation (**Supplemental Results 2.2, Fig. S3**). Our first approach was to determine whether it can report similar results to those obtained using H_2_^18^O_2_ with apo-PcuC. To do so, we designed a slightly different strategy as chemical probes are usually used at several concentrations to identify reactive residues in activity-based protein profiling or related approaches [36,38]. Thus, we incubated the apo-PcuC with 0 to 10 molar equivalents of probe and quantified the labeling of each Met after trypsin proteolysis and LC-MS/MS (**Fig. 2C, E**). The overall pattern closely matched the one obtained with H_2_^18^O_2_: Met25 was unlabeled, while Met135 was the most reactive, reaching ∼83% with 10 equivalents of probe. Both Met88 and Met128 were found labeled around 5%. Similarly, Met132 labeling percentage (∼44%) was nearly identical to the values obtained with H_2_^18^O_2_. Met88 and Met128 showed low labeling (∼5%), and Met132 reached ∼44%, consistent with H_2_^18^O_2_ results. Differences were observed for Met63 and Met90, that were less labeled (∼20% and ∼11%, respectively), whereas Met61 showed higher labeling (∼51%). These discrepancies could arise from differences between the H_2_^18^O_2_ and oxaziridine protocols (kinetics versus concentration range) or from the slightly larger size and aromatic moiety of the oxaziridine probe, which may influence interactions with residues surrounding Met differently than peroxide. At the highest probe concentration, the reactivity ranking of Met residues in apo-PcuC was: Met25, Met88, Met128, Met90, Met63, Met132, Met61, Met135. Consistently with the results obtained with H_2_^18^O_2_, this shows that copper-coordinating Met63 and Met88 do not present a sensitivity to oxidation particularly different from that of the other residues. These results indicated that the use of the oxaziridine probe is relevant to quantify Met sensitivity to oxidation in copper-loaded PcuC. To our knowledge, although it was previously used to identify oxidizable Met residues [36,39], the reactivity of the oxaziridine probes was never compared to physiological oxidants. This showed that, despite few differences, the oxaziridine probe effectively mimics H_2_O_2_-induced oxidation and is very likely relevant to identify redox-active Met residues in proteins from various biological contexts.

### 2.3. Cu binding strongly alters PcuC’s Met sensitivity to oxidation

We then sought to evaluate whether the presence of copper in PcuC affects Met sensitivity to oxidation using the same approach. First, we incubated the protein with increasing concentration of CuSO_4_ and observed that two coppers could be bound to the protein using non-denaturing ESI-MS (**Fig. S4A**), as shown for the *B. diazoefficiens* homolog [24]. As anticipated, the incubation of the copper-loaded PcuC with 1,000 equivalents of H_2_^18^O_2_ resulted in a quick and complete protein degradation (**Fig. S4B**). This was very likely due to Fenton-like reaction occurring between the peroxide and the copper, either bound to the protein or free in the solution because of remaining trace despite the desalting. Using the probe on the copper-loaded PcuC revealed a very different profile than for the apo protein (**Fig. 2D, E**). Indeed, we first noticed that all Met residues were labeled, including Met25. By comparison with the apo-PcuC, we observed strong increases for Met25, Met63, Met88 and Met90 labeling. The most labeled became Met61, followed by Met132, while the level of labeling of Met135 strongly decreased. Interestingly, the reactivity of the copper-ligand Met63 and Met88 increased by 2.1-fold and 3.5-fold, respectively (**Fig. 2D, E**). This indicates that copper-binding increases Met sensitivity to oxidation. At the highest probe concentration, the reactivity ranking in Cu-loaded PcuC was: Met128, Met25, Met88, Met135, Met63, Met90, Met132, and Met61. The change in reactivity of the copper-coordinating Met63 and Met88 could be attributed to alterations in the electronic density on sulfur atoms, as copper coordination withdraws electron density, increasing the electrophilicity of sulfur. Additionally, copper binding might stabilize the Met lateral chain in a conformation that favors oxidation. The increased sensitivity of the other Met not directly coordinating copper, may result from the initial oxidation of the coordinating Met63 and Met88, which could alter the protein’s folding, thereby enhancing the overall accessibility and reactivity of other Met residues. Such initial oxidation events has been shown to increase protein hydrophobicity and potentially favor further oxidation [40,41]. The apparent decrease in the sensitivity of Met135 is most likely due to the increase in sensitivity of the other Met residues, as it is unlikely that the structural properties have been modified for this Met (the C-terminal part remained unfolded, *vide infra*).

### 2.4. PcuC coordinates two Cu ions at the canonical Cu binding site

To evaluate whether structural determinants can explain the observed differences of Met residues reactivity, we solved the tridimensional structures of the apo and copper-bound PcuC by X-ray crystallography (**Fig. 3**). The structure of the apo-PcuC was solved at 1.05 Å resolution in a *P*4_3_2_1_2 space group obtained from a crystal grown in sodium citrate at pH 8.5 (**Table S1**). The structure is typical for homologs of this protein family, i.e. a seven stranded β-barrel with one side of this barrel containing three additional strands, which include the conserved Met and His residues involved in copper binding. Two parts of the protein could not be resolved: the first spans residues Ala55 to Met61, while the second corresponds to the C-terminus from Arg124 and includes Met128, Met132 and Met135 (**Fig. 3A**). These parts are very likely flexible in the protein, and the C-terminal part was predicted to be disordered (**Fig. S5**). To obtain experimental data on copper binding to PcuC, we added ten copper equivalents to the protein and re-screened for crystallization. We obtained crystals that clearly showed two sites of high electron density, very likely corresponding to metals, but they were affected by translational non-crystallographic symmetry (**Supplemental Results 2.4**). Then, by mixing the copper-loaded PcuC with 20% PEG 6000, 1 M LiCl_2_ and 0.1 M citric acid at pH 4.0 in a second crystallization attempt, we obtained crystals that diffracted to 2.75 Å resolution and the elucidated structure showed the presence of two copper ions (**Table S1, Fig. 3B**). One copper (Cu1) is bound to Met88 Sδ (2.6 Å) and His86 Nδ (2.1 Å), similarly to what is observed in the PcuC homologs from *Streptomyces lividans* [42] and *Thermus thermophilus* [23]. However, in these homologs the copper is coordinated by two other ligands (corresponding to His51 and Met63), whereas in the structure of *C. sphaeroides* PcuC a water molecule linked the copper to Nδ of His51 (2.7 Å; Cu1-HOH = 3 Å), and Met63 adopted such a conformation that its side chain was not in the coordination sphere of Cu1 (the distance Cu1 to Met63 Sδ is 5.5Å). Finally, what we interpreted as a citrate molecule, very likely coming from the crystallization media, participated in the coordination sphere of this copper with a bidentate interaction from one carboxylate moiety (Cu1-OA1 = 2.4 Å and Cu1-OA2 = 2.9Å). Overall, although the position of this copper binding site is conserved between the three homologs structurally characterized, the coordination sphere is different in *C. sphaeroides* PcuC. The additional copper ion (Cu2) observed in the structure of *C. sphaeroides* PcuC was bound by Nε of His53 (1.8 Å) and three oxygen atoms from the three carboxylate moieties of citric acid (2.7 Å, 3.1Å and 3.1 Å; **Fig. 3B**). Altogether, these results show that PcuC could bind two coppers without the involvement of the unfolded C-terminal part. This finding was unexpected given the results obtained for the *B. diazoefficiens* PcuC homolog [24]. While it is possible that the two chaperones function differently, several elements support a unified model that reconciles both sets of results. In this model, a first copper ion is stably coordinated in the canonical site (high-affinity), while a second copper ion oscillates between various positions along the C-terminal extension and the coordination site observed in our structure (low-affinity) (**Supplemental Discussion**).

**Fig. 3.**
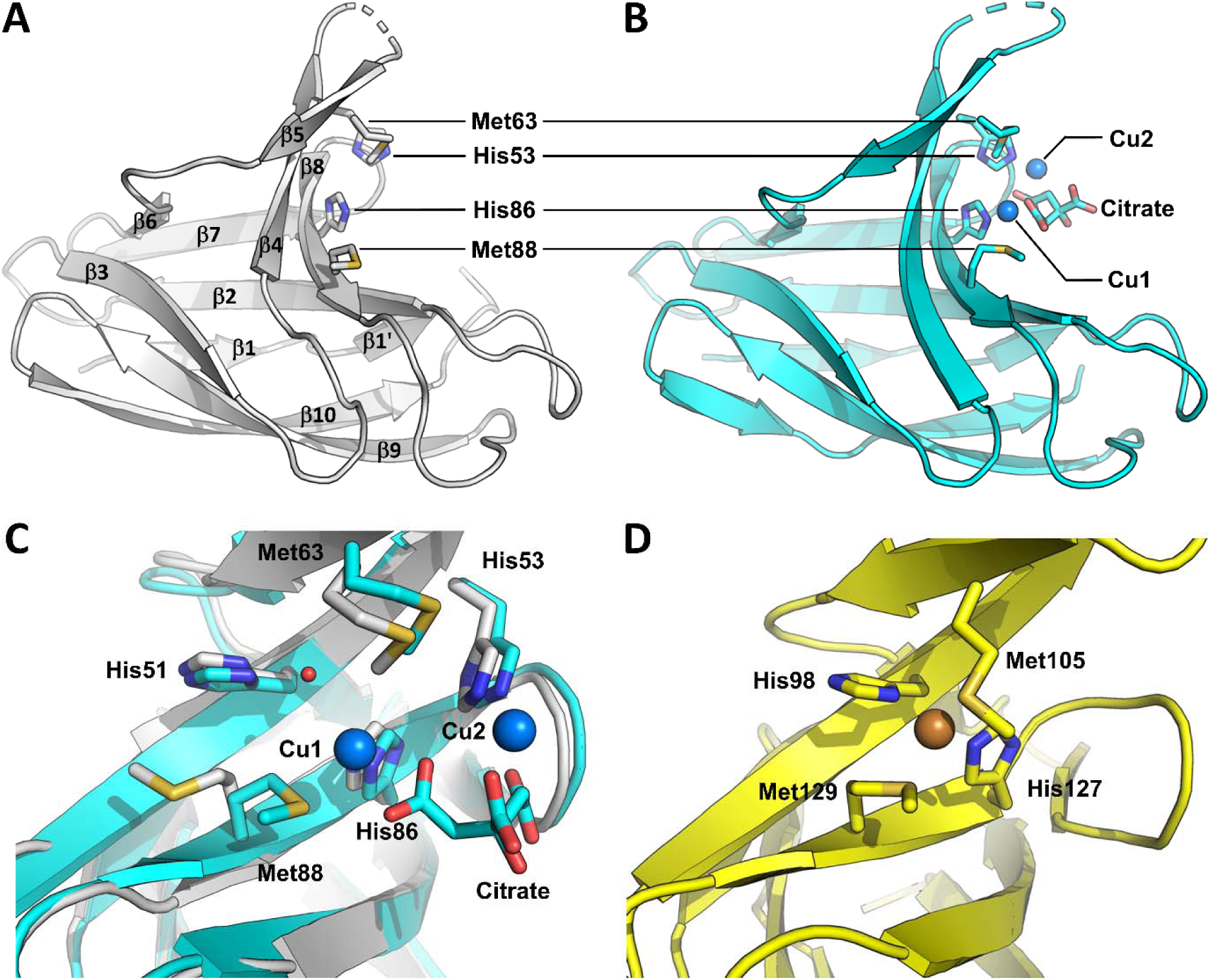
Structural characterization of apo-PcuC and Cu-loaded PcuC. Overall structure of the apo form of PcuC (PDB 9REZ) **(A)** and the Cu-loaded form of PcuC (PDB 9RF0) **(B)** highlighting the copper-coordinating residues Met63, Met88, His53, and His86, as well as the citrate molecule, shown as sticks; copper ions are shown as *blue* spheres. **(C)** Superimposition of both the apo-and copper-bound form of PcuC with the side chains of Met63, Met88, Met90, Met132, and Met135 shown as sticks. **(D)** Details of the Cu-binding site of *S. lividans* PcuC (PDB 3ZJA) showing Met105, Met129, His98, and His127 coordinating copper, which is shown as an orange sphere.

### 2.5. Met oxidation sensitivity correlates with solvent exposure in the folded region but with positive charge in the unfolded C-terminal extension

To understand the factors influencing Met oxidation in PcuC, we analyzed the solvent exposure and local charges surrounding each Met in both apo- and Cu-bound structures. As the unfolded C-terminal extension containing Met128, Met132, and Met135 was unresolved, we used an AlphaFold model to complete the analysis (**Fig. 4A**). Overall, the model fitted very well with the resolved structure of the apo-PcuC, with a root mean square deviation (RMSD) of only 0.390 Å. Met25 to Met90 are located at very similar positions in both the resolved structure and the predicted model, and only slight differences exist for the lateral chain positions of Met61, Met63, Met88 and Met90. Moreover, on the model, the C-terminal extension containing Met128, Met132 and Met135 was depicted without secondary structure and with a low confidence score, as expected for a disordered region. In the folded region (encompassing Met25 to Met90), solvent exposure values were consistent between the resolved and modeled structures. Met25 showed minimal exposure, while Met61 and Met63 were among the most exposed. Met128, Met132, and Met135, in the unfolded C-terminal extension, displayed high solvent exposure in the model (**Fig. 4B**). We assumed that the C-terminal extension remains unstructured in both the apo- and Cu-PcuC, and the exposure values for these Met residues are similar to the model. To determine whether oxidation sensitivity correlates with solvent exposure, we plotted the values of oxidation determined with ^18^O-labeled hydrogen peroxide and oxaziridine as function of the Met exposure values for both the apo-and the copper-loaded PcuC structures (**Fig. 4C, D**). In apo-PcuC, oxidation levels correlated positively with solvent exposure for Met residues in the folded region (R² = 0.48 for H_2_^18^O_2_ and R² = 0.90 for oxaziridine). A similar positive correlation was observed using the solvent exposure values from the AlphaFold model (**Fig. S6**). Surprisingly, the C-terminal Met residues showed a strong negative correlation between oxidation and solvent exposure (R² = 0.99). Similar trends were observed in Cu-loaded PcuC, though with lower correlation values (**Fig. 4C**). Strikingly, the most exposed Met128 was among the least oxidized, both with H_2_^18^O_2_ and the oxaziridine probe, and regardless of copper presence. To explain this discrepancy in the disordered region, we considered the electrostatic environment. Surface charge maps revealed that, while most of the protein surface is neutral or negatively charged, the C-terminal region includes a localized positive charge due to a lysine (**Fig. 4E, F, G**). We quantified the partial charge within 5.5 Å of each Met sulfur (**Fig. 4H**). In the folded domain, no correlation was found between oxidation and local charge (**Fig. S7A, B**). However, for the C-terminal Met residues, we observed a strong positive correlation between oxidation sensitivity and positive local charge (R² = 0.92–0.97 in apo-PcuC) (**Fig. 4I**). These results indicated that the oxidation sensitivity of the Met residues belonging to the unfolded C-terminal part of PcuC was positively correlated with the positive partial charge surrounding the sulfur atom of the Met. Together, these results reveal that distinct factors govern metal oxidation in different structural contexts of PcuC: solvent exposure is crucial in the folded part, while local positive charge controls reactivity in the unfolded part. These results are consistent with the fact that solvent exposure plays very likely a major role in Met sensitivity to oxidation, as shown previously [43–45], but that other factors should be considered [46,47]. This suggests that Met sensitivity to oxidation might be protein and Met dependent and should be determined for each protein of interest.

**Fig. 4.**
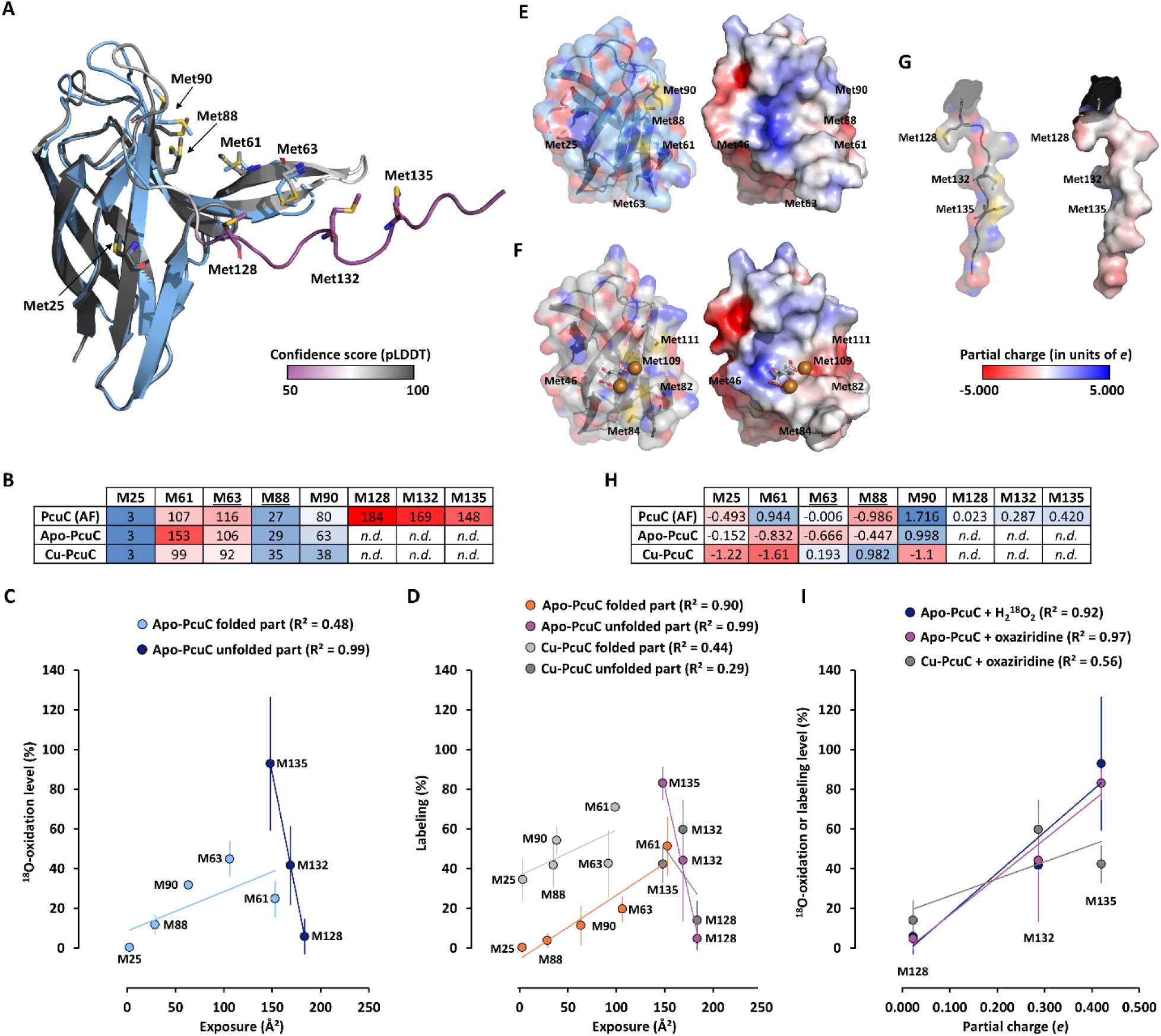
Correlation of Met oxidation sensitivity with solvent exposure and local charge. **(A)** Met position in resolved apo-PcuC structure (*light blue*) superimposed with AlphaFold 2.0 model. RMSD = 0.390 Å for 627 atoms. The model is colored according to predicted local distance difference test (pLDDT) values (pLDDT < 70: low confidence (*purple*); pLDDT > 70 (*white*): confident; pLDDT > 90: high confidence (*dark grey*)). Mean pLDDT = 84.69. Met are shown in stick with carbon colored as the main color of the proteins. Sulfur, oxygen and nitrogen are *yellow*, *red* and *blue*, respectively. **(B)** Heatmap showing the solvent exposure (Å²) of the Met, using a *blue*-to-*red* color scale, with *blue* and *red* representing the lowest and highest exposure, respectively. *n.d., not determined.* Underlined Met63 and Met88 belong to the Cu-binding site. **(C)** Plot of oxidation level of the Met residues as function of solvent exposure for the apo-PcuC after 60 min oxidation with H_2_^18^O_2_ with independent linear regressions for the Met residues of the folded part and the unfolded C-terminal part. **(D)** Plot of oxidation level of the Met residues as function of solvent exposure for the apo- and Cu-loaded PcuC after exposure to 10 equivalents of oxaziridine with independent linear regressions for the Met residues of the folded part and the unfolded C-terminal part. **(E, F, G)** Visualization of surface colored by atom (*left part*, colored as in **(A)**) and partial electronic charge (*right part*) for the apo-, the Cu-loaded PcuC and the C-terminal part of the AlphaFold model. Citrate is shown in stick with carbon and oxygen in *grey* and *red*, respectively. Copper atoms are shown as *orange* spheres. **(H)** Heatmap showing the partial charge of the environment surrounding the sulfur of the Met, using a *blue*-to-*red* color scale, with *blue* and *red* representing the more negative and positive charge, respectively. *n.r., not resolved.* Underlined Met63 and Met88 belong to the canonical Cu-binding site. **(I)** Plot of oxidation of the Met residues from the C-terminal unfolded part determined with H_2_^18^O_2_ and oxaziridine as function of the partial charge of the environment surrounding the sulfur of the Met determined on the AlphaFold model for the apo- and Cu-loaded PcuC with linear regressions.

## 3. Conclusion

In this study, we investigated Met oxidation sensitivity in the copper chaperone PcuC from *C. sphaeroides* using an oxaziridine-based probe and H_2_^18^O_2_. Our results show that the probe reproduces Met oxidation patterns obtained with hydrogen peroxide, supporting its use as a reliable tool for quantitatively mapping Met oxidation sensitivity in proteins, including metalloproteins. Met63 and Met88, which contribute to the copper-binding site, do not display enhanced oxidation sensitivity in the absence of copper; however, copper binding markedly increases their susceptibility to oxidation, highlighting a direct interplay between metal coordination and local Met reactivity. A limitation of our study is that it relies exclusively on *in vitro* assays and therefore does not capture the full complexity of the cellular environment, including copper homeostasis and potential interactions between PcuC and other periplasmic proteins. *In vivo*, oxidative conditions are expected to differ, with low concentrations of hydrogen peroxide but the potential for locally generated, copper-driven hydroxyl radicals through Fenton-like chemistry. Such conditions could alter oxidation patterns and potentially affect residues beyond Met. Our results also provide new insights into the function of the conserved, disordered C-terminal extension present in most PcuC homologs. This region may play a dual role by facilitating copper incorporation into the coordination site and protecting against oxidative damage via the highly oxidation-sensitive Met135. Moreover, its location within an unfolded region of PcuC is expected to favor its repair, as Msr, and particularly the periplasmic MsrP, display enhanced activity toward oxidized Met residues in unfolded proteins [20,41]. Together, these features suggest that Met135 may act as a reversible redox buffer, for which cyclic oxidation and reduction could protect the protein to support efficient copper delivery to CoxB and bacterial respiration under copper stress. To investigate this hypothesis *in vivo*, future studies could assess whether a strain expressing a PcuC variant in which Met135 is replaced by a non-oxidizable residue such as leucine retains the ability to support CoxB assembly and O□ production across a range of copper concentrations. In parallel, performing these analyses in a strain lacking MsrP would help evaluate the contribution of Met repair to PcuC function under copper stress. Such a dual role in copper insertion and protection against oxidation could be envisioned for other copper transporters and oxidases containing Met-rich regions [17,48]. Beyond the case of PcuC, our findings suggest that copper-binding could render a protein more prone to oxidation by generating localized ROS and enhancing the oxidation susceptibility of nearby Met residues. Such properties could potentially occur in other copper-binding proteins, including the prion protein and α-synuclein, for which copper redox chemistry contributes to oxidation and protein dysfunction in disease contexts. In this broader framework, the oxaziridine-based strategy provides a versatile chemical biology platform for proteome-wide mapping of Met oxidation susceptibility across diverse organisms and physiological conditions. Applied broadly, this approach could identify oxidation-sensitive Met hotspots in metalloproteins and redox-regulated pathways, with potential applications in redox signaling studies, biomarker discovery, and the identification of redox vulnerabilities that may be therapeutically exploited in pathogens or disease-associated proteins.

## 4. Methods

### 4.1. Protein and chemicals

PcuC (Uniprot# Q3J4W6) was produced and purified as described [49]. ^18^O-labeled hydrogen peroxide and dibenzocyclooctyne-Cyanine5 fluorophore (DBCO-Cy5) were purchased from Sigma-Aldrich. Initial concentration of commercial H_2_^18^O_2_ was estimated to be ∼450 mM using FOX method [50]. The oxaziridine probe ((4-azidopiperidin-1-yl)(3-phenyl-1,2-oxaziridin-2-yl)methanone) was synthetized according to [37] and validated by NMR (**Fig. S8**). Protein concentrations were determined using the Bradford method (Bio-Rad Protein Assay Dye Reagent Concentrate). SDS-PAGE were performed using Bolt^TM^ pre-cast gel system (ThermoFisher) using 12% polyacrylamide gels and 2-(N-Morpholino)ethanesulfonic acid (MES) running buffer. Visible and fluorescent imaging were performed using Amersham ImageQuant™ 800 (Cytiva).

### 4.2. PcuC oxidation, Cu-loading and labeling with the oxaziridine probe

Apo-PcuC (70 µM) was oxidized with 10, 100, 500 or 1,000 molar equivalents (equiv.) of H_2_O_2_ in 50 mM 3-(N-Morpholino)propanesulfonic acid (MOPS) buffer at pH 8.0 for 1 h and the reaction was stopped by adding excess dithiothreitol (DTT) (100 mM) and subsequent desalting using Sephadex G-10 (PD MidiTrap G-10, Cytiva). For oxidation with H_2_^18^O_2_, PcuC (100 µM), apo or Cu-loaded, was incubated for 2, 5, 15, 30, 45 and 60 min with 1,000 equiv. of H_2_^18^O_2_ in 50 mM MOPS, pH 8.0, then the reaction was stopped with the addition of 0.071 U.µl^-1^ of catalase followed by desalting. PcuC Cu-loading was performed by incubating the protein (200 µM) with 2 to 10 equiv. of CuSO_4_ in MOPS 50 mM followed by desalting. For Cu-loading before oxaziridine labeling, 5 equiv. of CuSO_4_ were used. Oxaziridine labeling was made by incubating PcuC (14 or 100 µM) with 0, 0.1, 0.5, 1, 5 or 10 equiv. of the probe for 20 or 30 min in 50 mM MOPS, pH 8.0 followed by desalting. For in-gel fluorescence analysis, oxaziridine-labeled PcuC was incubated with 10 equiv. of DBCO-Cy5 overnight in 50 mM MOPS, pH 8.0 and excess reagent was removed by ultrafiltration (5 kDa cutoff, Vivaspin 500, Sartorius).

### 4.3. Mass spectrometry analyses

ESI-MS analyses were performed as described [20]. For proteolysis and LC-MS/MS analyses, 20 µL of PcuC solution (7 µM) in 50 mM MOPS buffer (pH 8.0) with 20% acetonitrile (ACN) were digested by adding 3 µL of trypsin/Lys C (Promega) at a concentration of 12.5 ng.µL^-1^ in 0.1% formic acid (FA) for 3 h at 37°C. For TMT labeling, the manufacturer’s protocol (ThermoFisher) was followed. Peptides from each experimental condition (various concentrations of oxaziridine or different reaction times with H_2_^18^O_2_) were incubated with one of the six 6-plex TMT reagents, consisting of reporter ions 126, 127, 128, 129, 130, and 131 (0.8 mg in 41 µL of anhydrous ACN) for 1 h at room temperature. The reactions were quenched by adding 2 µL of 5% ammonium hydroxide for 15 minutes. For each experiment (oxaziridine or H_2_^18^O_2_), the six conditions were pooled and diluted in 300 µL of 0.1% trifluoroacetic acid (TFA). The labeled peptides were filtered using a C18 column (ThermoFisher) to remove excess TMT reagents, eluted with 2×300 µL of 50% ACN/0.1% TFA, and dried using a SpeedVac. The dried samples were reconstituted in 2% ACN/0.1% FA to achieve a theoretical protein concentration of 1 µg.µL^-1^. The samples were injected into an LC-MS/MS system using a high-performance nano-liquid chromatography (UHPLC RSLC, ThermoFisher) equipped with online desalting (PEPMAP C18, 5 µm, 300 Å trap column, 300 µm × 5 mm), coupled to a Q-Exactive Plus hybrid mass spectrometer (ThermoFisher). The analytical column used was an Acclaim PepMap RSLC capillary column (nanoViper C18, 2 µm, 100 Å, 75 µm × 15 cm). The gradient for peptide elution was as follows: solvent A: 0.1% FA, solvent B: ACN with 0.1% FA. The gradient program included the following steps: (1) 2.5–25% B over 57 min, (2) 25–50% B over 6 min, (3) 50–90% B in 1 min, and (4) 90% B for 10 min. A data-dependent acquisition (DDA) method was employed with a Top10 approach, where the 10 most abundant peptides were selected for fragmentation in each survey scan. MS settings were: precursor mass range 375 to 1400 m/z, resolution 70,000, AGC target 3×10^6^, maximum IT 100 ms. MS/MS parameters: fragment mass range 100 to 2000 m/z, resolution 70,000, AGC target 2×10^5^, and normalized collision energy of 27 (33 for TMT-labeled samples). Peak list files in .mgf format (Mascot Generic Format) were generated using Proteome Discoverer 2.4 (ThermoFisher). Protein identification was carried out using MASCOT 2.3 software, querying a database containing PcuC and common contaminants (246 sequences, including trypsin, keratins, etc.). Search parameters were set as follows: trypsin as the digestion enzyme, up to 4 missed cleavages, precursor mass tolerance of 10 ppm, product ion mass accuracy of 0.02 Da, with fixed modifications of TMT6plex (N-term) and TMT6plex (K), and variable modifications including deamidation (Gln, Asn), ^16^O-oxidation (Met, monoisotopic mass = 15.994915 Da), ^18^O-oxidation (Met, monoisotopic mass = 17.99916), ^16^O-dioxidation (Met, monoisotopic mass = 31.989829 Da), ^18^O-dioxidation (Met, monoisotopic mass = 35.998321 Da), ^16^O/^18^O-dioxidation (Met, monoisotopic mass = 33.994075 Da) or oxaziridine modification (Met, monoisotopic mass = 167.080710 Da). Quantification was performed by exporting reporter ion intensity data for all peptides to Excel using Mascot software. The percentages were determined by summing, for each Met, the relative intensities of all detected forms. For the H_2_^18^O_2_ treatment, ^18^O-containing oxidation states (^18^O-sulfoxide, ^18^O-sulfone, and ^16^O/^18^O-sulfone) were grouped as oxidized Met, while non-modified and ^16^O-oxidized forms, considered artifactual, were counted as non-oxidized. Note that, according to the manufacturer, the ^18^O-labeled hydrogen peroxide solution may contain up to 10% of H_2_^16^O_2_ but because the exact value was not determined, this parameter was not considered for calculation. Similarly, for the oxaziridine treatment, non-modified and ^16^O-oxidized forms were considered as non-labeled.

### 4.4. PcuC crystallization and structure analysis

Crystals of PcuC were obtained by the sitting drop method using 96 wells plates and a pipetting robot (TECAN Evoware). The apo protein was concentrated to 26 mg.mL^-1^ and crystals were obtained by mixing 1 µL of protein with 1 µL of 1.2 M sodium citrate and 0.1 M Tris pH 8.5. Crystals quickly appeared and, after supplementing the mother liquor with 15% glycerol, a few crystals were flash frozen directly in liquid nitrogen. Crystals were collected at the SOLEIL synchrotron (France) and the structure of apo PcuC was then solved at 1.05 Å resolution in a *P*4_3_2_1_2 space group. To obtain experimental data on copper binding to PcuC, ten copper equivalents were added to the protein and crystallization of the protein was rescreened. Crystals were obtained in a condition containing 25% PEG 3350 and 20 mM CaCl_2_, 20 mM CdCl_2_ and 20 mM CoCl_2_. This crystal form diffracted to 2.1 Å resolution, however it is affected by a translational NCS (tNCS) that, together with the three molecules in the asymmetric unit, appeared to be difficult to refine (*R* = 30.2%; *R*_free_ = 36.5%). This structure is therefore not presented, although it clearly showed three sites of high electron density, all of them relatively close to each other. Data collection at different wavelengths (copper and cobalt edges) also indicated that one site was occupied by copper, whereas the two others were cadmium ions. Additional screens were used to find a different crystal form that would not grow in a complex mixture of metal and that would eventually not be affected by tNCS. A crystal form was obtained in a condition containing 20% PEG 6000, 1 M LiCl_2_ and 0.1 M citric acid at pH 4.0. It diffracted to 2.75Å resolution and was refined to a *R* and *R*_free_ of 24.2% and 30.4%, respectively.

Prediction of intrinsically disordered region was made using IUpred2A [51]. Solvent exposure of Met was determined using the “*get_area*” function with default parameters and partial charges were determined using APBS Electrostatics plugin of PyMol 3.1.0a0 (open-source build).

## Supplementary data

Supplemental Results, Supplemental Discussion, Fig. S1-S8, Table S1, Supplemental References.

## Data availability

The mass spectrometry proteomics data have been deposited to the ProteomeXchange Consortium via the PRIDE [52,53] partner repository with the dataset identifier PXD062219. Structure data have been deposited in Protein Data Bank (PDB) with accession code 9REZ for the apo form and 9RF0 for the copper-bound form.

## Acknowledgments

The authors kindly thank the support team of Aix Marseille Univ., INRAE, Biodiversité et Biotechnologies Fongiques – Chantal Parodi-Negri, Naura Thibeau and Christophe Boyer. This research was funded by the Commissariat à l’Energie Atomique et aux Energies Alternatives (CEA) and by the project METOXIC [ANR 16-CE11-0012]. This work received support from the French government under the France 2030 investment plan, as part of the Initiative d’Excellence d’Aix-Marseille Université — IM2B (AMX-19-IET-006) for L.M. and L.T. – A*MIDEX (AMX-21-PEP-028) for L.T, and from the NovoNordisk foundation (OxyMiST project, grant number NNF20OC0059697).

## CRediT author statement

**Lionel Tarrago:** Conceptualization, Methodology, Validation, Formal analysis, Investigation, Resources, Data Curation, Writing - Original Draft, Writing - Review & Editing, Visualization, Supervision, Funding acquisition, Project administration. **Lise Molinelli:** Conceptualization, Methodology, Validation, Formal analysis, Investigation, Data Curation, Writing - Review & Editing. **Maya Belghazi:** Conceptualization, Methodology, Validation, Formal analysis, Investigation, Data Curation, Writing - Review & Editing. **Mathilde Tribout:** Formal analysis, Investigation, Data Curation. **David Lemaire:** Methodology, Validation, Formal analysis, Investigation, Data Curation. **Pierre Legrand:** Methodology, Validation, Formal analysis, Investigation, Data Curation. **Sandrine Grosse:** Formal analysis, Investigation, Data Curation. **David Pignol**: Conceptualization, Methodology, Supervision, Funding acquisition, Writing - Review & Editing. **Monique Sabaty:** Conceptualization, Methodology, Supervision, Funding acquisition, Writing - Review & Editing. **Thierry Tron:** Conceptualization, Methodology, Supervision, Funding acquisition, Writing - Review & Editing. **Pascal Arnoux:** Conceptualization, Methodology, Validation, Formal analysis, Investigation, Resources, Data Curation, Writing - Original Draft, Writing - Review & Editing, Visualization, Supervision, Funding acquisition, Project administration.

## Declaration of generative AI in scientific writing

During the preparation of this work the authors used ChatGPT to improve the clarity and readability of the text. After using this tool, the author reviewed and edited the content as needed and take full responsibility for the content of the publication

