## Supplementary data for "Quantitative mapping of methionine sensitivity to oxidation in the copper-bound PcuC chaperone"

### Supplementary Results

#### 2.1. PcuC's methionine residues are oxidizable by $H_2O_2$ and repairable by MsrP

To determine whether Met residues can be oxidized, we first incubated the purified recombinant protein with increasing concentrations of non-labeled hydrogen peroxide (0 to 1,000 molar equivalents (equiv.)). We observed delays of migration of the treated protein on polyacrylamide gel proportional to the concentration of oxidant (**Fig. S1A**). This delay indicates that Met are oxidized, as shown for other proteins (1–3). Measurement of the total protein mass by ESI-MS showed first that the untreated purified PcuC had a mass of 14,440 Da, corresponding to its expected mass, and thus was free of copper (**Fig. S1B**). Then, it showed that the oxidative treatment resulted in the formation of a heterogeneous population of oxidized protein with multiple mass increases of 16 Da, up to seven at the highest concentration of $H_2O_2$ (1,000 equiv.). To validate that the 16 Da increases correspond to the formation of MetO, we used the specific MetO reductase activity of MsrP (4). Upon treatment of the oxidized PcuC by MsrP, we observed a significant restoration of its electrophoretic mobility on SDS-PAGE (**Fig. S1C**). Moreover, MsrP was active only on the oxidized PcuC, with activity value equivalent to those obtained with efficiently reduced substrates (4) (**Fig. S1D**). Finally, ESI-MS analysis of oxidized PcuC after MsrP's treatment showed a diminution of the measured mass down to that of the native protein and with no more than three oxidized Met remaining (**Fig. S1E**). Altogether, these results indicate that the treatment of PcuC with $H_2O_2$ induces the formation of up to seven MetO and that MsrP can efficiently repair oxidized Met of PcuC.

#### 2.2. The oxaziridine probe binds the apo-PcuC

To evaluate whether the oxaziridine probe can bind to PcuC, we incubated the apo-protein with increasing amounts of probe. Then, taking advantage of the azide handle, we performed a click reaction with a dibenzocyclooctyne (DBCO) moiety conjugated to a cyanine-5 (Cy5) fluorophore. We revealed PcuC functionalization by in-gel fluorescence and observed that fluorescence intensity increased with the amount of probe used (**Fig. S3A**). Then, we analyzed the total mass of PcuC after reaction with the probe by Matrix-Assisted Laser Desorption/Ionization Time-of-Flight mass spectrometry (**Fig. S3B**). While the spectrum of the non-treated apo-PcuC contains a single peak corresponding to the theoretical mass of the protein, the spectrum corresponding to the protein treated with the oxaziridine displayed several overlapping peaks, with mass increases corresponding to one to five adducts of the probe on the protein. These results confirm the ability of the oxaziridine probe to label the protein, and its suitability to quantify Met sensitivity to oxidation.

#### 2.4. A preliminary structure of PcuC suggests it can bind two metal ions

To obtain experimental data on copper binding to PcuC, we added ten copper equivalents to the protein and re-screened the crystallization of the protein. We obtained crystals in a condition containing 25% PEG 3350 and 20 mM $CaCl_2$, 20 mM $CdCl_2$ and 20 mM $CoCl_2$. This crystal form diffracted to 2.1 Å resolution, however it is affected by a translational non-crystallographic symmetry that, together with the three molecules in the asymmetric unit, appeared to be difficult to refine ($R = 30.2\%$; $R_{free} = 36.5\%$). This structure is therefore not presented, although it clearly showed two sites of high electron density, both relatively close to each other. Data collection at different wavelengths (copper and cobalt edges) also indicated that one site was occupied by copper, whereas the other was a cadmium ion.

### Supplementary Discussion

#### A unified model for copper binding in PcuC integrates structural and NMR observations

The three-dimensional structure of the holo-PcuC revealed that two copper ions are coordinated in the canonical H53$x_n$M63$x_{22}$H86xM88 site and a citrate molecule, and that residues from the unfolded C-terminal part are not involved in the coordination (**Fig. 2**). These results were unexpected considering the data obtained for the *B. diazoefficiens* PcuC homolog (5). Nuclear magnetic resonance (NMR) analyzes have suggested that a Cu(I) ion can bind to the canonical site, while a Cu(II) ion can bind to the C-terminal region. This hypothesis was based on the fact that incubation with copper shifted the NMR signals of the C-terminal Met, both in the full-length protein and in a peptide corresponding to the C-terminal segment. Although the two chaperones may differ in their mechanisms, several lines of evidence support a unified model that can account for both sets of observations. In this model, a first copper ion is stably coordinated in the canonical site (high-affinity), while a second copper ion oscillates between various positions along the C-terminal extension and the coordination site observed in our structure (low-affinity). The C-terminal region would thus guide the copper ion to the coordination site, and our structure would be a snapshot picture of the protein with the second copper near the coordination site. The supporting arguments are:

1) The Met53 and His63 residues that coordinate the second copper in our structure are highly conserved in PcuC. In contrast, the C-terminal extension contains abundant Met and His residues, but these are not conserved in specific positions (5). This suggests that the C-terminal extension facilitates copper recruitment, similar to bacterial multicopper oxidase CueO (6) or the human copper transporter Ctr-1 (7).

2) ESI-MS analysis of both *B. diazoefficiens* (5) and *C. sphaeroides* PcuC revealed two populations of PcuC, each bound to either one or two copper atoms, suggesting that one copper is more weakly coordinated to the protein than the other.

3) Electron paramagnetic resonance analysis of *B. diazoefficiens* PcuC showed a wide distribution of distances between the two copper ions, with a full width at half maximum ranging from 1.6 to 2.6 nm. This indicates a dynamic distance between the two atoms, suggesting that the second copper ion oscillates along the C-terminal extension, or that the extension itself oscillates with the copper coordinated by specific Met/His residues. In our structure, the short distance between the two copper ions (5.4 Å / 0.54 nm) might be due to the stabilizing effect of the citrate in the crystallization buffer.

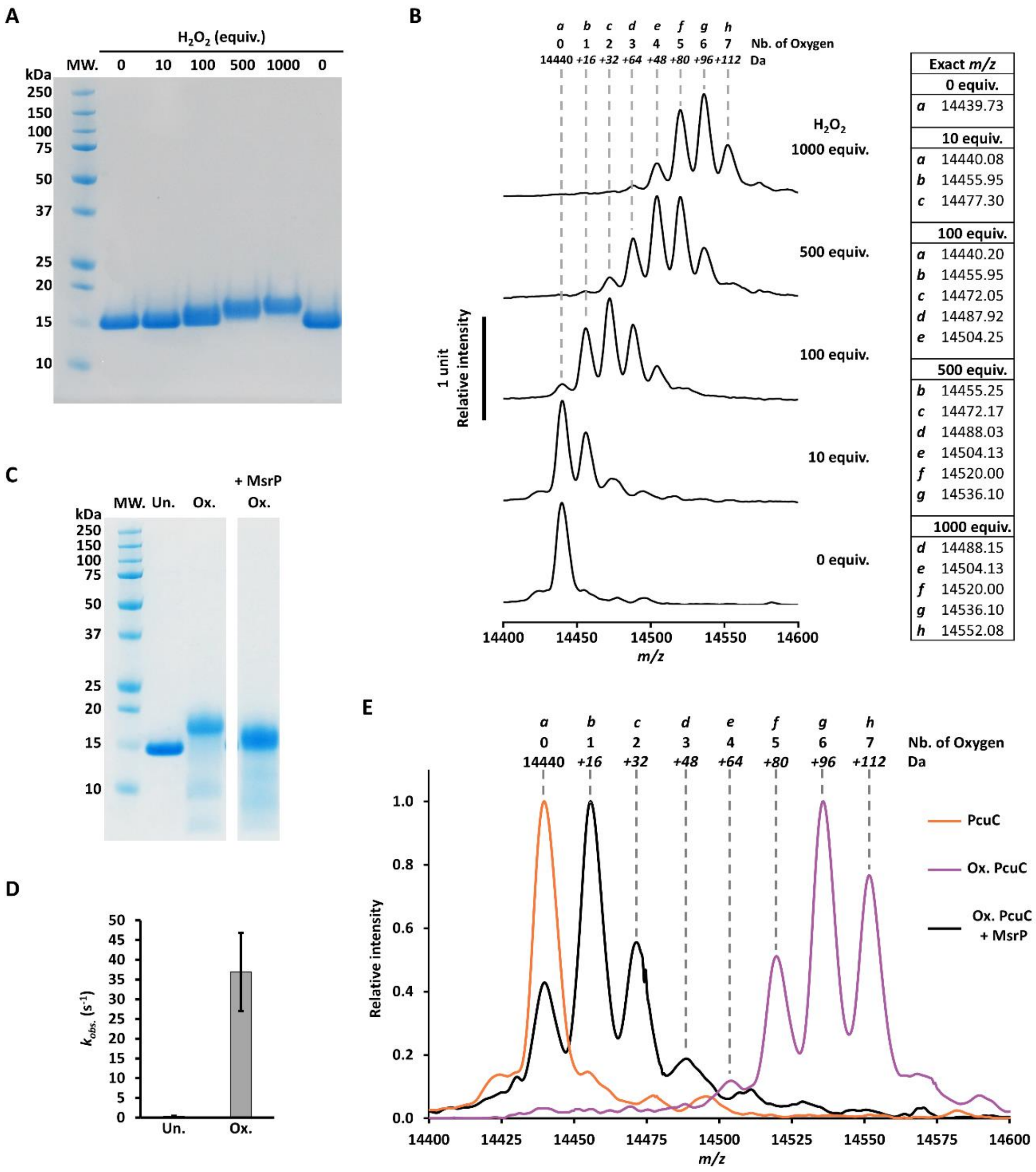

**Fig. S1. Oxidation of PcuC with hydrogen peroxide and reduction by MsrP.**

**(A)** Apo-PcuC (5 µg) was oxidized with 10, 100, 500 or 1,000 molar equivalents (equiv.) of hydrogen peroxide for 1 h and the reaction was stopped with excess DTT and subsequent desalting, then the samples were analyzed by SDS-PAGE. **(B)** Spectra of same samples analyzed by ESI-MS. **(C)** Migration profile of PcuC on polyacrylamide gel after oxidation and reduction by MsrP. PcuC oxidized with 1,000 equiv. of $H_2O_2$ was incubated with the MetO specific reductase MsrP (46 nM). After reaction, 5 µg were loaded on SDS-PAGE (*'+ MsrP Ox.'*) together with the unoxidized (*'Un.'*) and the oxidized protein (*'Ox.'*). **(D)** MsrP activity using oxidized PcuC as substrate. MsrP activity was measured using 50 µM of oxidized PcuC as substrate as described (4). **(E)** Determination of the total masses of untreated PcuC, oxidized PcuC and oxidized PcuC reduced by MsrP by ESI-MS. Exact values of *m/z*: *a*, 14439.73 (PcuC); *a,* 14439.85 (Ox. PcuC + MsrP); *b*, 14455.72; *c*, 14471.47; *d*, 14488.62; *e*, 14504.02; *f*, 14519.77; *g*, 14535.75; *h*, 14551.62. *MW., Molecular weight markers.*

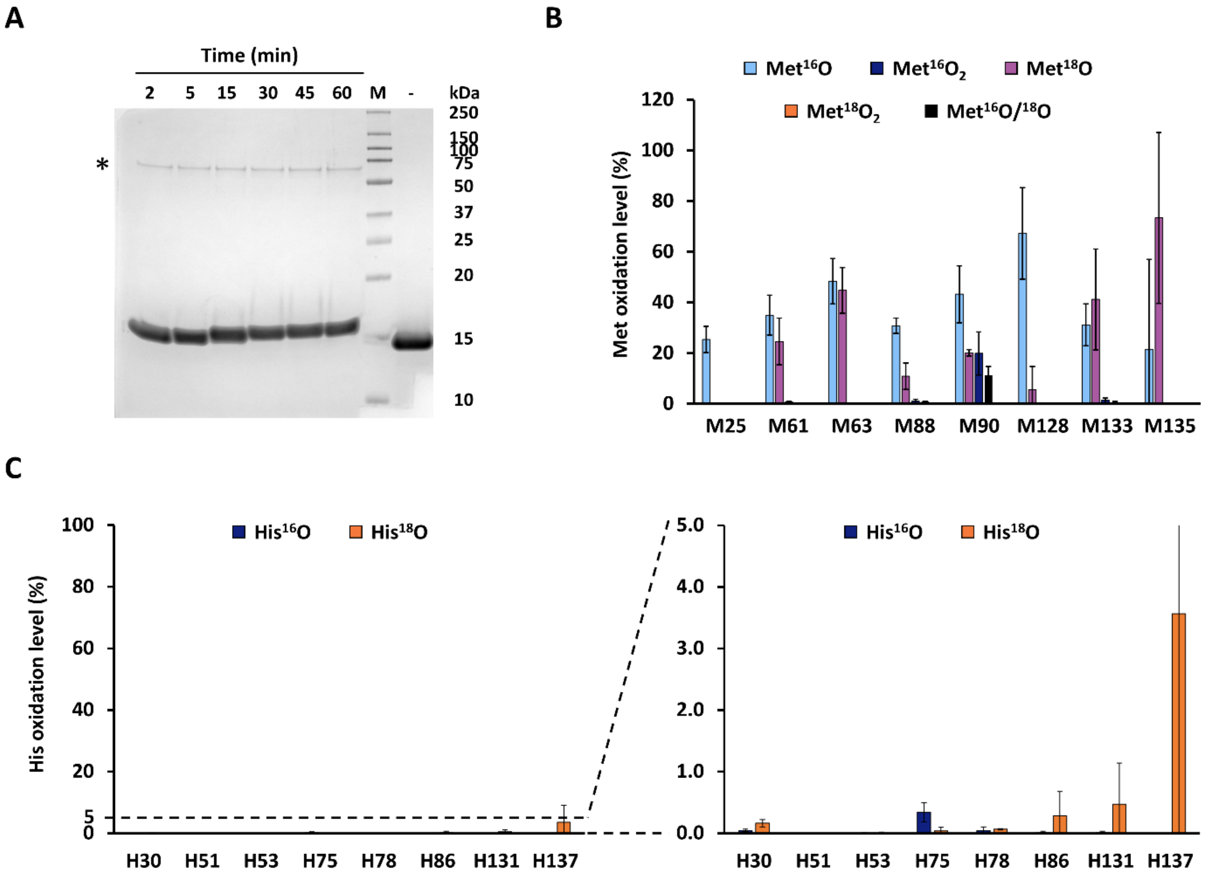

**Fig. S2. Oxidation of apo-PcuC with $H_2{}^{18}O_2$.**

**(A)** Representative migration profile of PcuC on polyacrylamide gel after kinetic oxidation with 1,000 equiv. of $H_2{}^{18}O_2$. Reaction was stopped after 2, 5, 15, 30, 45 and 60 min with catalase. Five µg were loaded on SDS-PAGE. The asterisk indicates the band corresponding to the catalase. The experiment was performed in triplicate and the samples were also analyzed by trypsin proteolysis and LC-MS/MS (*see main Fig. 1C*). **(B)** Percentage of the different forms of Met oxidized with $^{16}O$ and $^{18}O$ at the final point of the kinetics (60 min). The $^{16}O$ forms arose from artifactual oxidation during the analysis. **(C)** Percentage of His oxidized with $^{16}O$ and $^{18}O$ at the final point of the kinetics (60 min). LC-MS/MS data were analyzed to search for oxohistidine. The panel on the right shows an enlarged view, with the y-axis scaled from 0 to 5%.

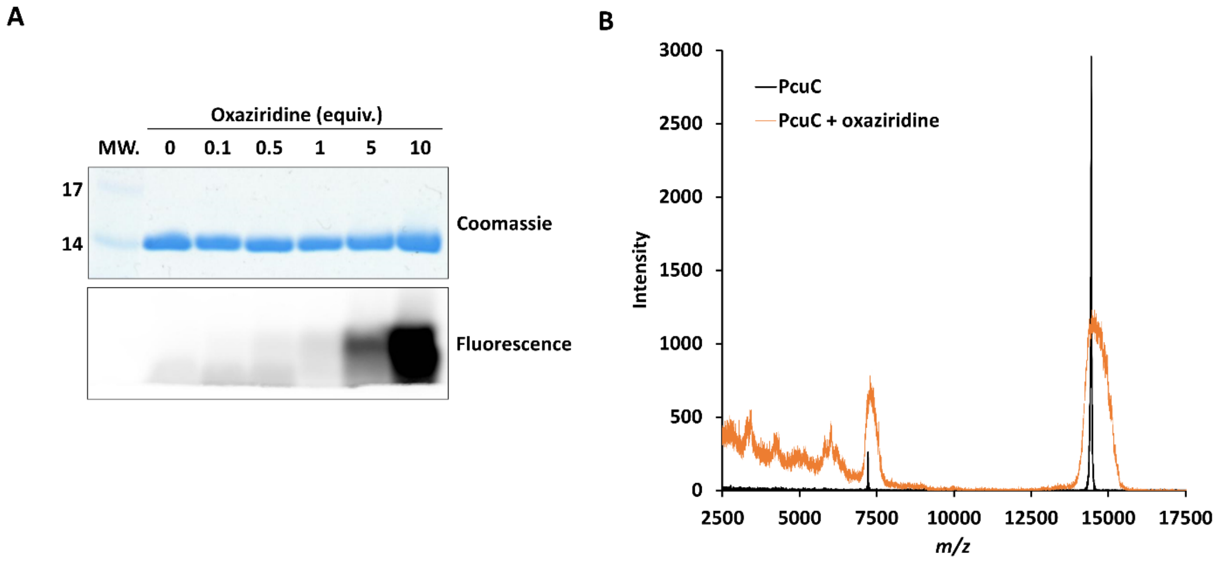

**Fig. S3. Reaction of the oxaziridine probe with PcuC.**

**(A)** SDS-PAGE analysis showing the reaction of PcuC with the oxaziridine probe, visualized through in-gel fluorescence. PcuC (14 µM) was incubated with increasing molecular equivalents (equiv.) of the oxaziridine probe for 30 minutes, followed by incubation with 10 equiv. of DBCO-Cy5 for 1 hour. Five µg of protein was loaded per lane and analyzed by SDS-PAGE. The gel was first analyzed for fluorescence ($\lambda_{excitation}$ = 635 nm; $\lambda_{emission}$ = 705 nm, bandwidth 40 nm), followed by Coomassie blue staining. **(C)** MALDI-TOF spectra of apo-PcuC before and after treatment with the oxaziridine probe. PcuC (100 µM) was incubated with 10 equiv. of the probe, desalted, and analyzed by MALDI-TOF.

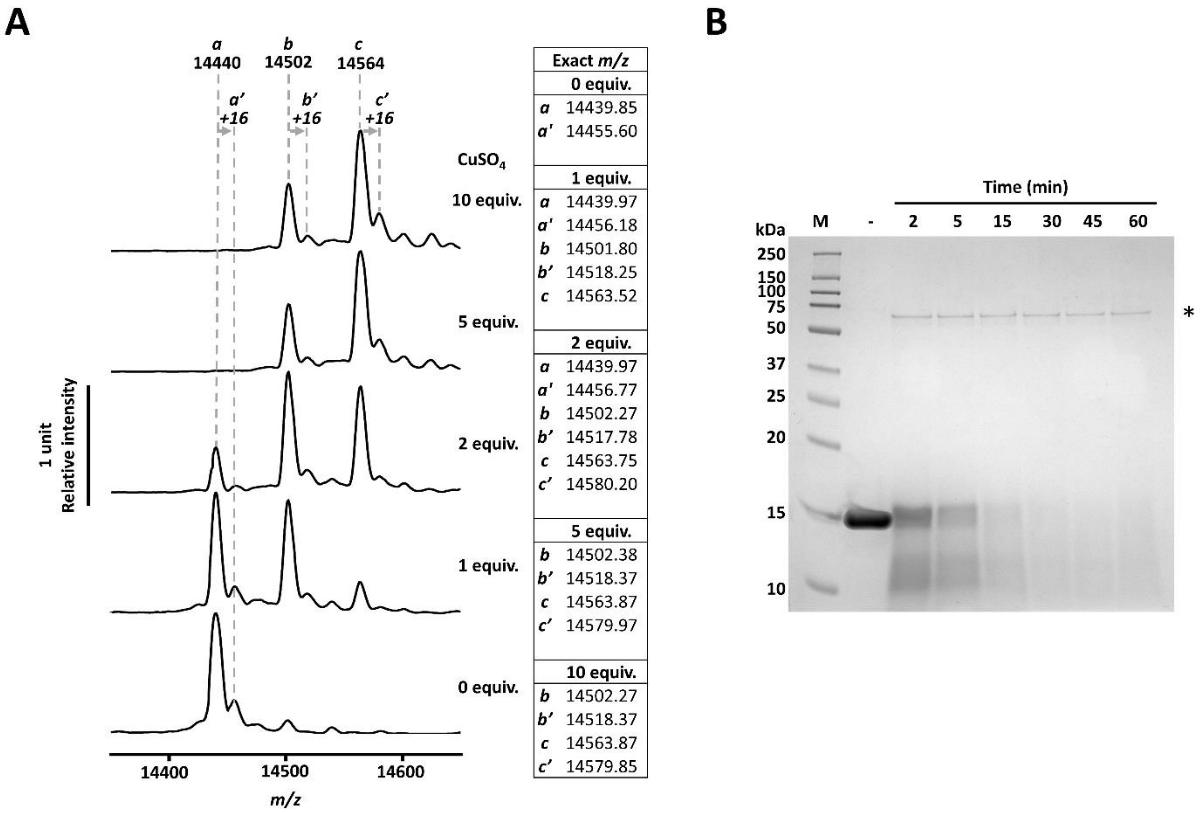

**Fig. S4. Copper loading in PcuC and effect of oxidation.**
**(A)** ESI-MS spectra of copper-loaded PcuC. Apo-PcuC (200 µM) was incubated with 0, 1, 2, 5 or 10 equiv. of $CuSO_4$ for 5 min and the solution was desalted, then the samples were analyzed by ESI-MS in non-denaturating condition. The peaks *a'*, *b'* and *c'* correspond to a small proportion of PcuC oxidized on one Met very likely. **(B)** Representative migration profile of copper-loaded PcuC (5 equiv. of $CuSO_4$) on polyacrylamide gel after kinetic oxidation with 1,000 equiv. of $H_2{}^{18}O_2$. Reaction was stopped after 2, 5, 15, 30, 45 and 60 min with catalase. Five µg were loaded on the gel. The asterisk indicates the band corresponding to the catalase. The experiment was performed in triplicate, but the samples were not analyzed by trypsin proteolysis and LC-MS/MS because of protein degradation (*see* **Fig. 1A**).

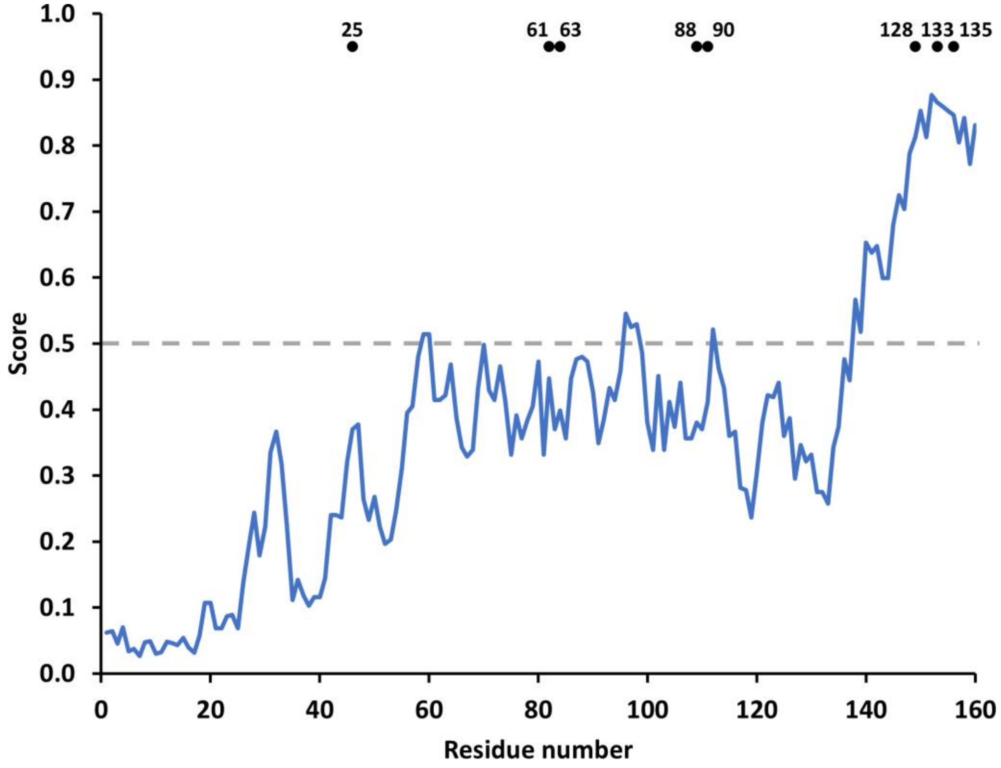

**Fig. S5. IUpred2A prediction scores for disordered regions in PcuC.**
The primary sequence of PcuC was analyzed using IUpred2A (8) to predict disordered regions. Each residue was assigned a score between 0 and 1, where a score of 1 represents the highest probability of disorder. Residues with a score above 0.5 are considered to be disordered. *Black* dots indicate the position of the eight Met.

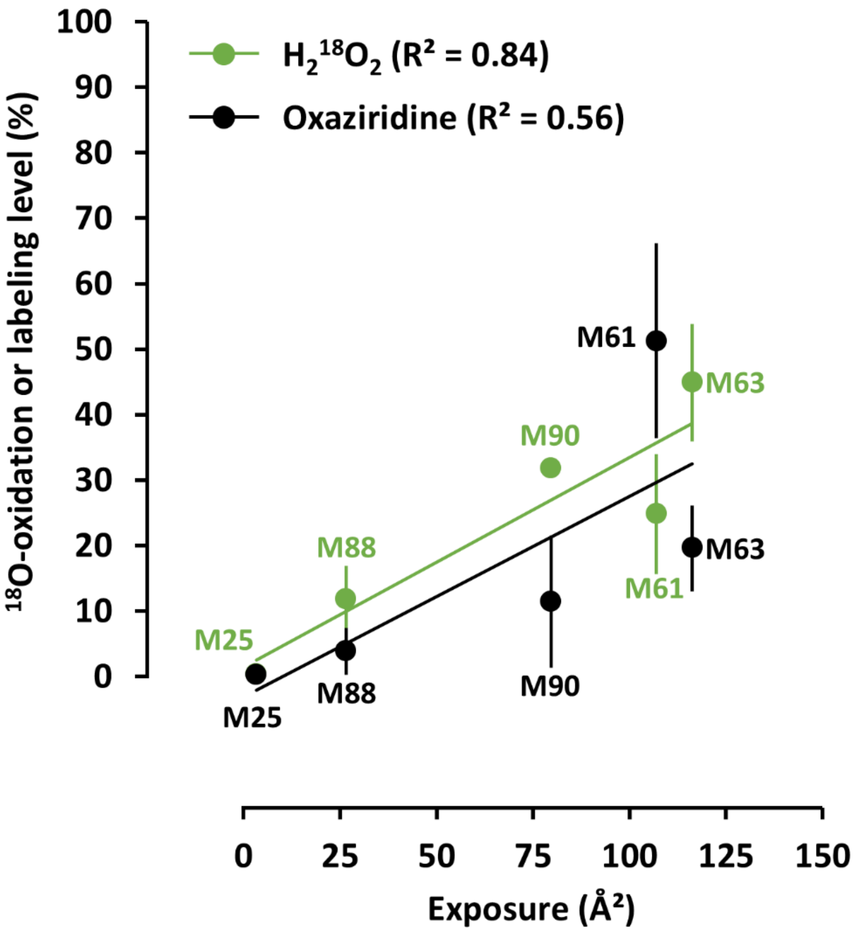

**Fig. S6. Correlation between solvent exposure on Met oxidation sensitivity in the AlphaFold model.** Plot of oxidation level of the Met from the folded part of PcuC obtained for the apo-PcuC after 60 min oxidation with $H_2{}^{18}O_2$ or 10 molar equivalents of oxaziridine as a function of solvent exposure determined for the AlphaFold model with linear regressions.

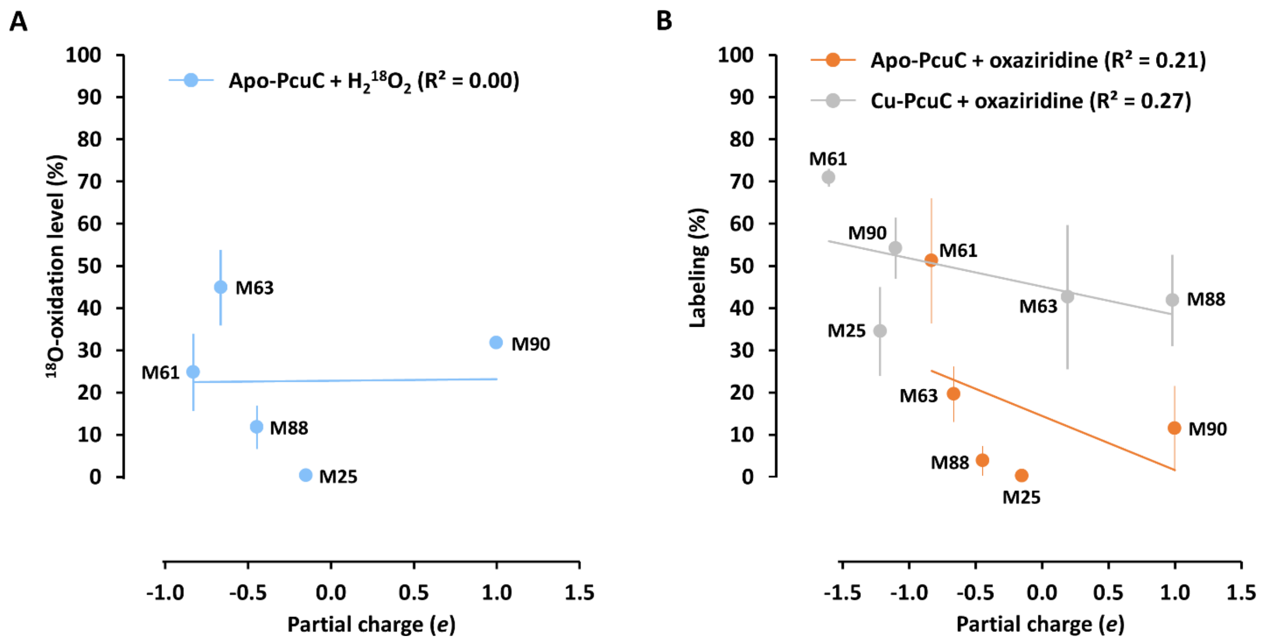

**Fig. S7. Partial charge of PcuC and oxidation levels of the Met from the folded part.**
**(A)** Plot of oxidation level of the Met as function of the partial charge of the environment surrounding the sulfur of the Met for the apo-PcuC after 60 min oxidation with $H_2{}^{18}O_2$ with linear regression. **(B)** Plot similar to (A) for the apo- and Cu-loaded PcuC after oxidation with 10 equiv. oxaziridine.

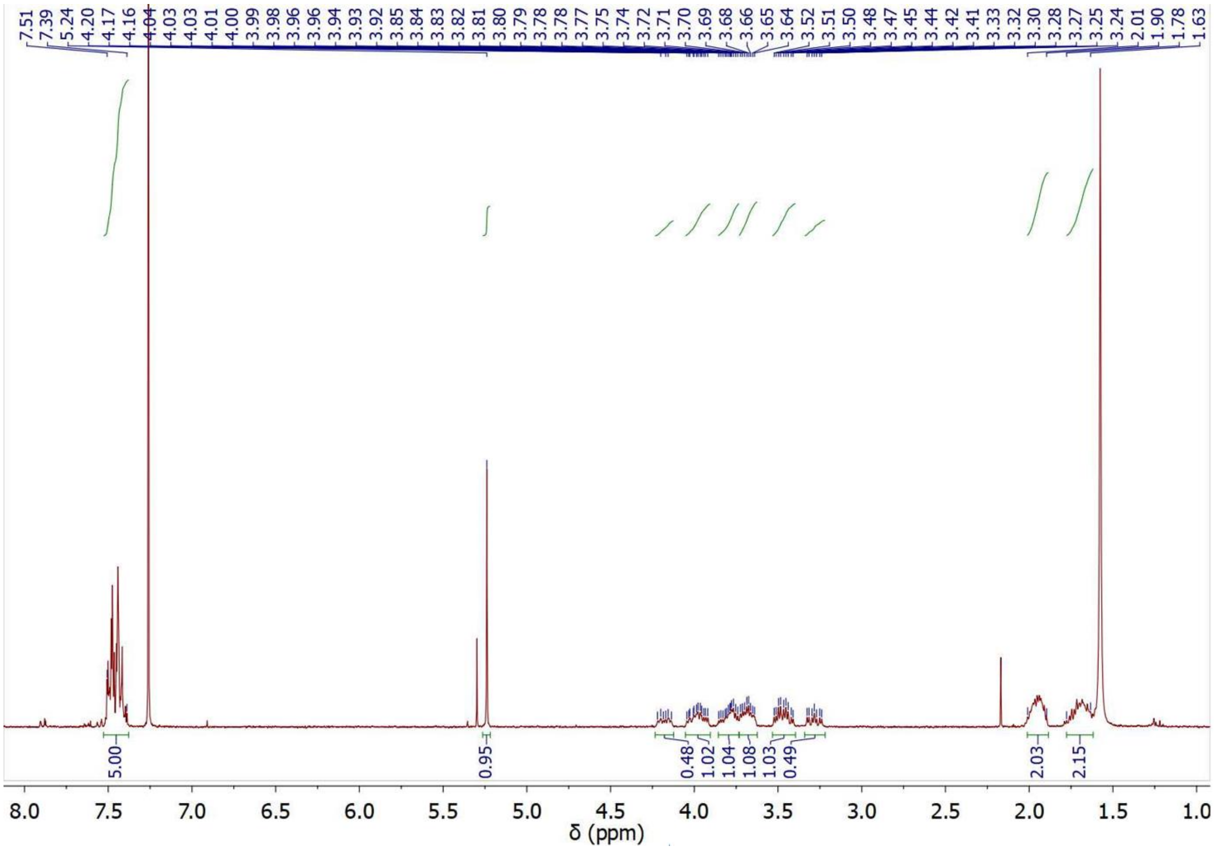

**Fig. S8. $^{1}H$ NMR spectrum of the oxaziridine probe**

**$^{1}H$ NMR:** (300 MHz, CDCl3) δ 7,51-7,39 (m, 5H), δ 5,24 (s, 1H), δ 4,24 – 4,13 (m, 1H), δ 4,06 – 3,91 (m, 1H), δ 3,87 – 3,74 (m, 1H), δ 3,68 (ddt, J = 10,6, 7,3, 3,5 Hz, 1H), δ 3,47 (ddt, J = 13,5, 9,7, 4,5 Hz, 1H), δ 3,28 (ddd, J = 13,3, 9,1, 3,5 Hz, 1H), δ 1,95 (m, 2H), δ 1,70 (m, 2H).

**Table S1. Crystal data, data-collection and refinement statistics.**

Values in parentheses correspond to the highest resolution shell.

#$R_{merge}$ = $S_{hkl}$ $S_i$ |$I_i(hkl)$ - <*I(hkl)*>| / $S_{hkl}$ $S_i$ $I_i$(hkl), where $I_i(hkl)$ is the *i*th observation of reflection *hkl* and <*I(hkl)*> is the weighted average intensity for all observations of reflection *hkl*.

| **Data collection** (PDB) | **9REZ** | **9RF0** |
|---|---|---|
| Data Set Name | Native | Cu |
| Wavelength | 0.918402 | 0.9786 |
| Beamline | PX1 / SOLEIL | PX1 / SOLEIL |
| Temperature | | 100K |
| Space Group | $P4_32_12$ | $P4_12_12$ |
| Unit-cell parameters (Å) | | |
| a | 48.1 | 55.5 |
| b | 48.1 | 55.5 |
| c | 110.6 | 81.7 |
| Resolution range (Å) | 50 – 1.05 | 50 – 2.75 |
| High resolution range (Å) | 1.11 – 1.05 | 2.9 – 2.75 |
| Observed reflections | 1,997,163 | 88,839 |
| No. of unique reflections | 61,261 | 3,690 |
| Completeness (%) | 99.5 (97.2) | 99.8 (98.5) |
| CC1/2 | 99.9 (73.0) | 99.8 (98.0) |
| <I/S(I)> | 18.1 (1.5) | 13.8 (2.6) |
| $R_{merge}$(%)# | 9.7 (163.2) | 19.1 (110.4) |
| **Refinement** | | |
| Resolution range (Å) | 44 – 1.05 | 46 – 2.75 |
| $R_{workt}$/ $R_{free}$ | 18.4 / 20.1 | 24.2 / 30.4 |
| No. of non-H atoms: | 1095 | 871 |
| Protein | 921 | 830 |
| Cu | - | 2 |
| Citrate | - | 13 |
| Water | 174 | 26 |
| B-Factors | | |
| Protein | 16.1 | 49.2 |
| Cu | - | 47.2 |
| Citrate | - | 83.2 |
| Waters | 29.8 | 42.3 |
| R.m.s. deviation from ideal geometry | | |
| Bond lengths (Å) | 0.013 | 0.007 |
| Bond angles (°) | 1.72 | 0.97 |
